# A Facile Method to Produce N-Terminally Truncated α-Synuclein

**DOI:** 10.1101/2022.02.21.481273

**Authors:** Rebecca J. Thrush, Devkee M. Vadukul, Francesco A. Aprile

## Abstract

α-Synuclein is a key protein of the nervous system, which regulates the release and recycling of neurotransmitters in the synapses. It is also involved in several neurodegenerative conditions, including Parkinson’s disease and Multiple System Atrophy, where it forms toxic aggregates. The N-terminus of α-synuclein is of particular interest as it has been linked to both the physiological and pathological functions of the protein and undergoes post-translational modification. One such modification, N-terminal truncation, affects the aggregation propensity of the protein *in vitro* and is also found in aggregates from patients’ brains. To date, our understanding of the role of this modification has been limited by the many challenges of introducing biologically relevant N-terminal truncations with no overhanging starting methionine. Here, we present a method to produce N-terminally truncated variants of α-synuclein that do not carry extra terminal residues. We show that our method can generate highly pure protein to facilitate the study of this modification and its role in physiology and disease. Thanks to this method, we have determined that the first six residues of α-synuclein play an important role in the formation of the amyloids.

## Introduction

α-Synuclein (α-Syn) is a 14 kDa intrinsically disordered protein that predominantly localizes at the presynaptic terminals of neurons [1, 2]. There, it is normally responsible for the recycling of neurotransmitter vesicles, mediating neurotransmitter storage and release [3-5]. In Parkinson’s disease (PD) and several other forms of neurodegeneration, this protein forms characteristic fibrillar aggregates, the amyloids, that are the major components of proteinaceous inclusions in the brain called Lewy Bodies (LBs) [6, 7].

α-Syn comprises of three domains: an amphipathic N-terminal tract with α-helical propensity (residues 1-60), an aggregation-prone central region called Non-Amyloid-β Component (NAC) that forms the core of the amyloid fibrils (residues 61-95), and a negatively charged C-terminal region that protects from aggregation and binds to metal ions, such as calcium (residues 96-140) [8, 9]. The N-terminal tract is a crucial region of α-syn, regulating several important functions of the protein [3-5, 10-12]. In fact, this domain is able to interact with lipid membranes, which, under normal conditions, regulates the fusion of neurotransmitter vesicles to the presynaptic button [5, 13]. In neurodegeneration, this interaction triggers the amyloid aggregation of the protein [10, 14]. Furthermore, it has been proposed that the N-terminal tract can establish long-range electrostatic interactions with the C-terminal region of α-syn [15-18], leading to more compact protein conformations [15, 19] and shielding the NAC region from aggregation [18]. It has also been proposed that alterations to these interactions may be driving α-syn liquid-liquid phase separation (LLPS) [20-22].

α-Syn undergoes several post-translational modifications (PTMs) that affect both its normal and pathological functions [23]. In particular, the protein is N-terminally truncated into fragments of different lengths *in vitro* [24, 25] and *in vivo* [26-29]. Some of these fragments have been reported to have different aggregation mechanisms and form structurally polymorphic amyloids with respect to the wild-type (WT) protein [11]. Despite the biological relevance of α-syn N-terminal truncations, investigations into these PTMs have been limited by the challenges associated with their synthesis. In fact, currently, most N-terminally truncated α-syn variants are produced with an additional starting Met [12, 30]. The presence of the extra Met allows for facile recombinant expression in *Escherichia coli* (*E. coli*). On the other hand, it may affect the binding to membranes and aggregation mechanisms of these proteins, which depend on just a few (∼10-20) N-terminal residues [11, 13, 31, 32]. To date, the only native N-terminal truncations to be recombinantly produced are those starting with a Gly at positions 14, 36 and 41 and have been generated with an *ad hoc* protocol based on placing the Gly downstream of the cleavage site for the tobacco etch virus (TEV) [11]. To expand the repertoire of α-syn N-terminal truncations that can be produced recombinantly, here, we described an intein-based protocol [33] to generate N-terminally truncated variants of α-syn with no overhanging residues. A key advantage of our method is that it does not require any proteases (e.g., TEV), making it time effective. As a proof-of-principle, we show the efficacy of our method in producing the N-terminally truncated variant of α-syn that lacks the first 6 residues (7-140 α-syn) [24].

## Materials and methods

### Fusion protein expression and purification

The intein-7-140 α-syn fusion protein cDNA was synthesized by GenScript and inserted into a pT7-7 plasmid. This plasmid will hereafter be referred to as pT7-7 Int7-140αSyn. BL21-Gold (DE3) competent *E. coli* cells (Agilent Technologies, USA) were transformed with the pT7-7 Int7-140αSyn plasmid, and the cells grown in 2 L of Overnight Express Instant LB medium (Merck, Germany) containing 100 μg/ml ampicillin at 28 °C for approximately 32 h. The cells were harvested by centrifugation and resuspended in 100 ml of ice-cold column buffer (20 mM 4-(2-hydroxyethyl)-1-piperazineethanesulfonic acid (HEPES), 0.5 M NaCl, 1 mM ethylenediaminetetraacetic acid (EDTA), pH 8.5) supplemented with EDTA-free protease inhibitors (Roche, Switzerland). 0.2 % Tween 20 was also added to reduce non-specific protein binding to the chitin resin. Cell lysis was induced by sonication on ice. Following centrifugation and syringe filtration, gravity chromatography was used to cleave and purify 7-140 α-syn [33]. To do so, the cell lysate was loaded on a chitin column equilibrated in column buffer. Unbound proteins were removed by washing the resin with 5 column volumes (CVs) of column buffer. The column was then equilibrated with 3 CVs of cleavage buffer (20 mM HEPES, 0.5 M NaCl, 1 mM EDTA, 50 mM β-mercaptoethanol (BME), pH 8.5) before the flow was stopped and the column incubated at room temperature for 44 h to facilitate cleavage. The target protein was eluted in 2 ml fractions using the column buffer. Fractions expected to contain α-syn (determined by absorbance at 275 nm using ε_275 nm_ = 5600 M^-1^cm^-1^) were combined and dialyzed overnight in PBS, pH 7.4 to remove the maltose binding protein (MBP) fragment and excess BME. The dialyzed sample was purified further by size-exclusion chromatography (SEC) using a HiLoad 16/600 Superdex 75 pg column (GE Healthcare, UK). Final protein concentration was determined by absorbance at 275 nm using ε_275 nm_ = 5600 M^-1^cm^-1^ as measured by UV-Vis spectrophotometry. Samples were taken throughout expression and purification for analysis by sodium dodecyl sulphate–polyacrylamide gel electrophoresis (SDS-PAGE) to determine the success and efficiency of each step.

### *Expression and purification of WT* α-syn

pT7-7 WT α-syn construct (Addgene, USA, gifted from Hilal Lashuel [34]), was transformed into BL21-Gold (DE3) competent *E. coli* cells and WT α-syn expressed according to the manufacturer’s instructions. Expression was scaled up to 1 - 5 L as desired and carried out at 28 °C overnight. The cells were harvested by centrifugation and resuspended in 20 mM Tris-HCl, pH 8.0 including 1 mM EDTA and protease inhibitors. WT α-syn was then purified as previously described [35].

### Electrospray ionization mass spectrometry

Purified protein samples (∼ 20 μM) in HPLC grade water were analyzed by electrospray ionization mass spectrometry (ESI-MS) to confirm molecular weight and sample purity. ESI-MS was performed by Lisa Haigh using the Chemistry Mass Spectrometry facilities available at the Molecular Sciences Research Hub, Department of Chemistry, Imperial College London.

### Beaded aggregation assay

50 μM monomer solutions were aggregated in the presence of 20 μM Thioflavin T (ThT) and 0.02% NaN_3_. 170 μl of each sample (5 replicates) was loaded into a 96 well full-area plate (non-binding, clear bottomed) and incubated at 37 °C for ∼ 140 h in a CLARIOstar Plus microplate reader (BMG Labtech, Germany). Aggregation was promoted through linear shaking (300 rpm, 300 s before each cycle) with the addition of a single borosilicate bead (3 mm diameter) to each well. Fluorescent intensity measurements were taken using spiral averaging (5 mm diameter) using excitation 440 nm, dichroic 460 nm and emission 480 nm filters, 3 gains and 50 flashes per well. Aggregation data was plotted in Prism and sigmoidal curves fit to the mean curve of each variant to estimate t_1/2_. t_lag_ was estimated as the x-intercept of the tangent at t_1/2_.

### Circular dichroism

The far-UV Circular Dichroism (CD) spectra of monomer samples (∼ 20 μM) were taken using a Jasco J-715 (Jasco Applied Sciences, Canada). Spectra were taken between 190 and 250 nm, scanning speed 50 nm min^-1^ and 15 accumulations. A background spectrum of the sample buffer was subtracted from both sample spectra. Raw data (units mdeg) was converted to mean residue ellipticity (MRE, units deg cm^2^ dmol^-1^) using [36]:

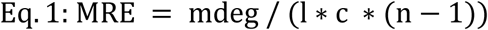

where mdeg is the raw data, n the number of amino acids, l the cuvette pathlength (mm) and c the protein concentration (M).

### Analysis of monomer conversion

The soluble and insoluble protein fractions after aggregation were separated by microcentrifugation (30 min, 16,900 g). Soluble fractions were then removed and analyzed by dot blot; 5 repeats of 5 μL aliquots were transferred to a nitrocellulose membrane which was then blocked before incubation with the anti-α-syn primary antibody MJFR1 (diluted 1:1000, abcam, UK). The membrane was subsequently incubated with the goat anti-rabbit IgG (H+L) highly cross-adsorbed secondary antibody conjugated to the fluorophore Alexa Fluor plus 555 (diluted 1:5000, Thermo Fisher Scientific, USA) before imaging. The insoluble fibril pellets were resuspended in fresh PBS, pH 7.4 and re-centrifuged to remove any remaining soluble protein. The washed fibrils were then resuspended in 250 μL of PBS, pH 7.4 and 10 μL applied to carbon-coated copper 300 mesh grids. Following negative staining with 2% (w/v) uranyl acetate the fibrils were imaged using the Tecnia 12 Spirit transmission electron microscope (Thermo Fisher Scientific (FEI), USA) available at the Electron Microscopy Centre, Center of Structural Biology, Imperial College London.

### Fibril digestion with proteinase K

20 μM solutions of the insoluble protein fractions after aggregation were incubated with increasing concentrations of proteinase K (0, 5, 15, 30 and 50 μg/ml) at 37 °C for 20 mins. The samples were separated by SDS-PAGE before transfer to a nitrocellulose membrane. The membrane was then blocked, incubated with primary and secondary antibodies and imaged as described above.

## Results and discussion

### Experimental design and model protein

As a model protein for our study, we chose the N-terminally truncated variant 7-140 α-syn due to its possible role as a precursor for shorter α-syn fragments [24]. Additionally, similar fragments, such as those comprising of the regions 5-140 and 10-122 of α-syn, have been found in patient brains [27, 28]. To express this protein, we designed a vector based on the *Saccharomyces cerevisiae*’s vacuolar ATPase (*Sce* VMA) intein [33]. Inteins are protein segments that are removed from within larger precursors by protein splicing, followed by chemical ligation of the flanking regions, which are called exteins. It has been shown that, when the intein is inserted within unrelated proteins, its splicing activity is retained provided that the first residue of the C-terminal extein is either a Cys, a Ser, or a Thr [37, 38]. Consequently, we engineered a construct, where the variant 7-140 α-syn was placed downstream of the *Sce* VMA intein. We also included a 10-residue fragment of the MBP at the N-terminus of the intein to facilitate expression, and the chitin binding domain (CBD) within the central region of the intein to enable affinity chromatography. It has been reported that insertion of the CBD does not affect the splicing activity of the VMA intein [33]. In our system, the intein splicing occurs in three steps while the protein is bound to a chitin resin (**Figure 1**): 1) the spontaneous N-S acyl rearrangement of the intein N-terminal Cys forms a thioester intermediate with the C-terminal residue of the MBP fragment, 2) we expose the protein to BME, which attacks the thioester intermediate resulting in N-terminal cleavage and removal of the MBP fragment, and 3) a spontaneous cyclisation at the C-terminal Asn takes place, which cleaves the truncated α-syn from the intein.

**Figure 1.**
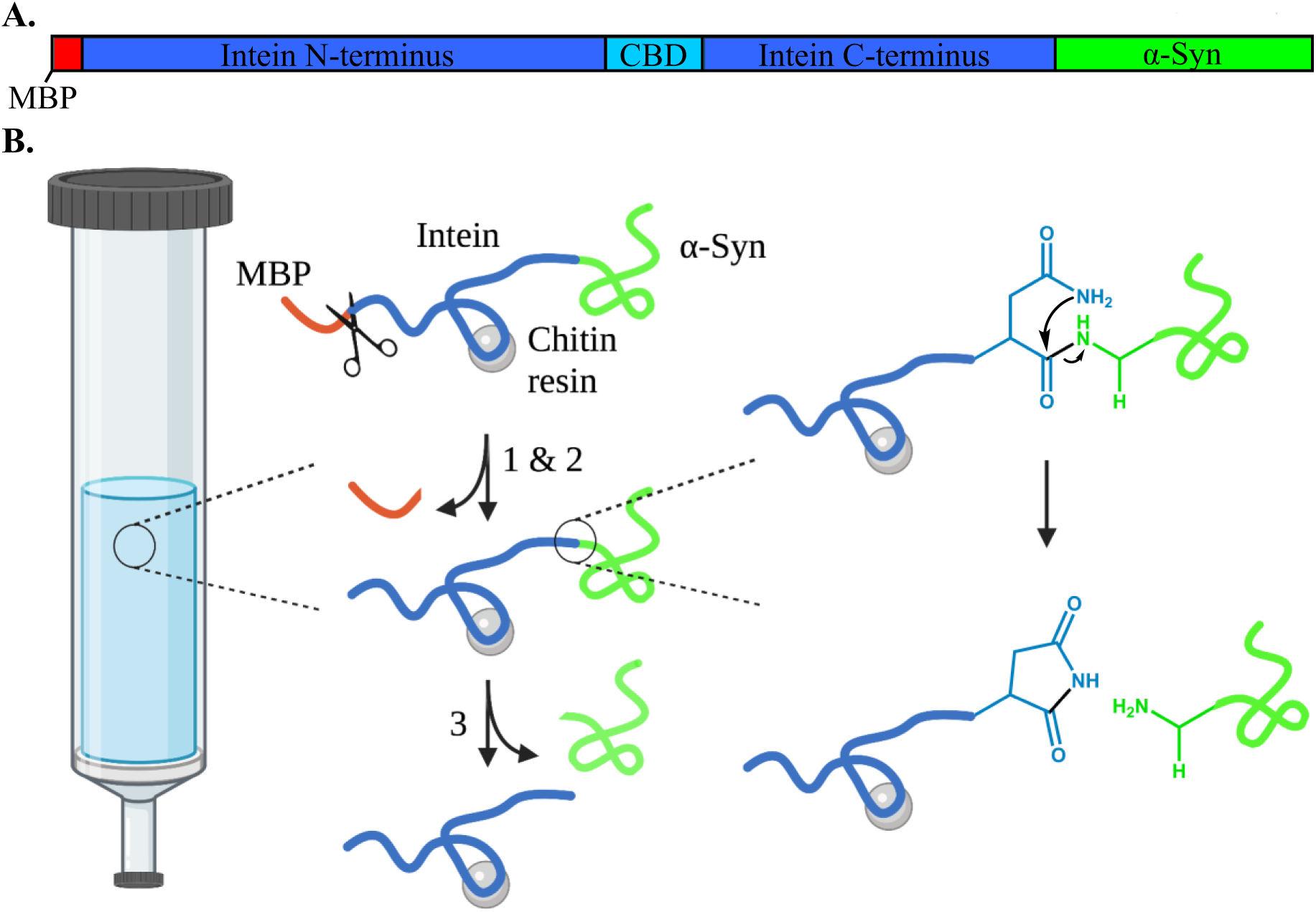
**A**. Schematic diagram depicting the domains of the intein-7-140 α-syn fusion protein. The MPB fragment is shown in red, the Sce VMA intein containing the CBD in blue and α-syn in green. This figure was created using SnapGene Viewer 6.0. **B**. representation of the VMA intein fusion protein cleavage reaction. 1) N-S acyl rearrangement at the intein N-terminus. 2) BME-mediated N-terminal cleavage. The BME is represented here as scissors. 3) spontaneous cyclisation at the intein C-terminus occurs, cleaving truncated α-syn from the protein. This figure was created using BioRender and ChemDraw 18.2.

### Expression and purification of 7-140 α-syn

To optimize the expression of the intein-7-140 α-syn fusion protein, we set up small scale cultures, where protein expression was induced with 1 mM IPTG at either 28 °C overnight or 37 °C for 4 h. Then, we analyzed the protein content of the cell extracts by SDS-PAGE. We found an enriched band of ∼ 70 kDa, which is the expected molecular weight of the fusion protein. This indicates that the protein is successfully expressed at both temperatures, with higher levels at 28 °C (**Figure S1**). Thus, we carried out our subsequent large-scale expression at 28 °C overnight. Following our optimization, we were able to obtain a yield of approximately 1.5 – 2 mg/L of cell culture of highly pure 7-140 α-syn.

The purity of the protein was confirmed by ESI-MS and SDS-PAGE (**Figure 2A and 2B**). We compared the CD spectrum of monomeric 7-140 α-syn to that of WT α-syn (**Figure 2C**). Both spectra show a high content in random coil, which is in agreement with the disordered conformation of the monomeric protein. We also analyzed the purified sample by native PAGE before and after boiling at 80 °C for 20 mins, mimicking the heat precipitation step used in WT α-syn purification (**Figure S2**). 7-140 α-Syn ran in a similar manner to the WT protein and boiling did not change the behavior of the truncated variant. Together this data show that our alternative purification method does not cause any significant alterations to the structural properties of the monomeric protein, relative to the protocol used for the WT protein.

**Figure 2.**
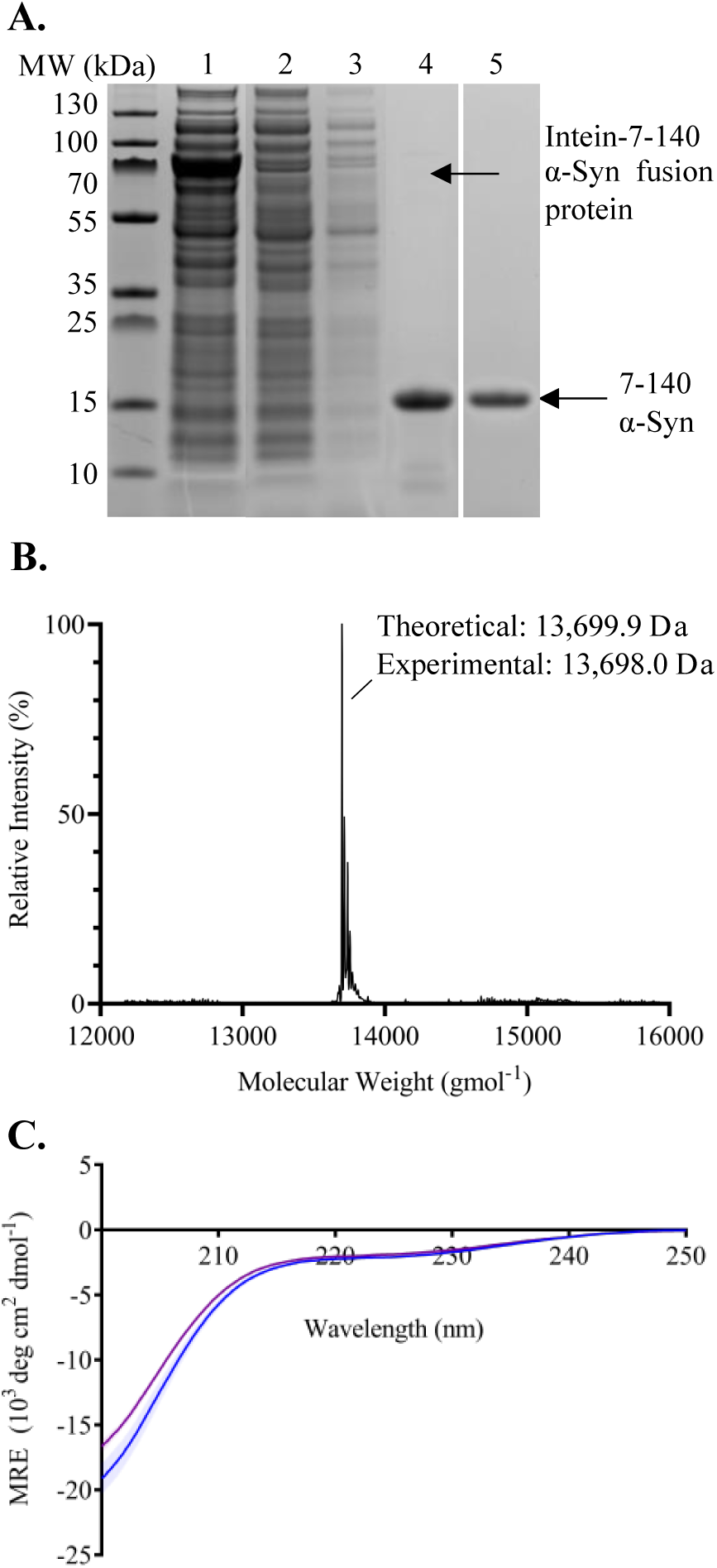
**A**. SDS-PAGE of samples taken throughout the purification process of 7-140 α-syn; crude extract (lane 1), crude extract after passing through column 5x (lane 2), flow through after washing the column with cleavage buffer (lane 5), compiled fractions predicted to contain the target protein after dialysis (lane 4) and the target protein purified by SEC (lane 5). **B**. The deconvoluted ESI-MS spectrum of 7-140 α-syn. Theoretical mass 13,699.9 Da, experimental mass 13,698.0 Da. **C**. The CD spectra of monomeric WT α-syn (blue) and 7-140 α-syn (purple). n = 3, error bars representing the standard deviation are shown as transparent bars.

### Characterization of the aggregation propensity of 7-140 α-syn

To characterize the aggregation of 7-140 α-syn, we performed ThT experiments, where 7-140 α-syn was incubated at 37°C under constant shaking in the presence of a borosilicate bead to accelerate aggregation. As a control, we analyzed WT α-syn under the same conditions (**Figure 3A**). We found that 7-140 α-syn aggregates at a significantly lower rate than WT α-syn. In particular, while the duration of the lag-phase (t_lag_) is the same for both proteins (∼30 h), the half-time of aggregation (t_1/2_) of 7-140 α-syn (69.7 ± 0.7 h) is approximately twice as long as WT α-syn (40.3 ± 4.8 h), suggesting that the growth phase of aggregation is affected by this truncation. Additionally, we observed that the ThT intensity at the plateau of aggregation of 7-140 α-syn is lower than that of WT α-syn (**Figure S3)**, suggesting that 7-140 α-syn generates a lower yield of amyloids than the WT protein. To validate this result, we performed a dot blot analysis to quantify the amount of soluble protein left at the plateau of aggregation. Our analysis revealed twice the percentage of soluble 7-140 α-syn remaining after aggregation compared to WT α-syn (**Figure 3B** and **S4**), confirming the lower fibril yield. This was further supported by transmission election microscopy (TEM) which showed that fibrils of 7-140 α-syn were overall shorter compared to WT α-syn (**Figure 4A** and **S5**). To determine if this increased fragmentation was associated to a lower fibril stability, the endpoints of aggregation were digested with proteinase K (**Figure 4B** and **4C**). We found that 7-140 α-syn fibrils are more susceptible to proteinase K digestion than the WT fibrils. In particular, we observed that the amount of undigested monomeric 7-140 α-syn decreases more rapidly than that of WT α-syn as a function of the protease concentration (**Figure 4C**).

**Figure 3.**
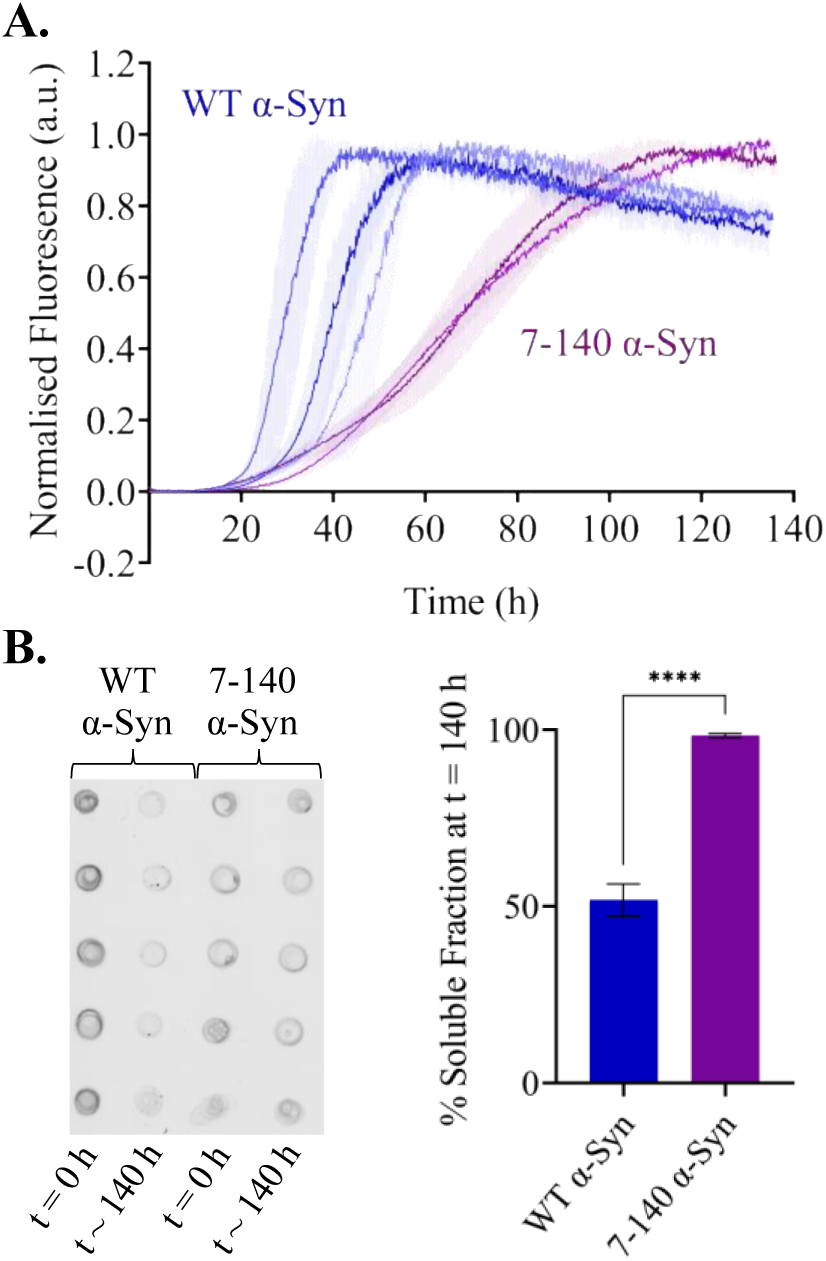
**A**. Beaded aggregations of WT (blue) and 7-140 (purple) α-syn. WT and 7-140 α-syn have t_1/2_ values of 37.2 ± 2.5 h and 69.7 ± 0.7 h and t_lag_ values of 28.4 ± 6.0 h and 30.7 ± 6.3 h respectively. The data shown is the mean of each normalized replica (n = 5, error bars are the standard deviation of the mean). **B**. Dot blot (left) of the soluble fraction before and after aggregation of WT and 7-140 α-syn for ∼ 140 h. The percentage of remaining soluble species (right) after aggregation for ∼ 140 h (n = 3, values represent means and error bars are the standard deviation, **** p < 0.0005).

**Figure 4.**
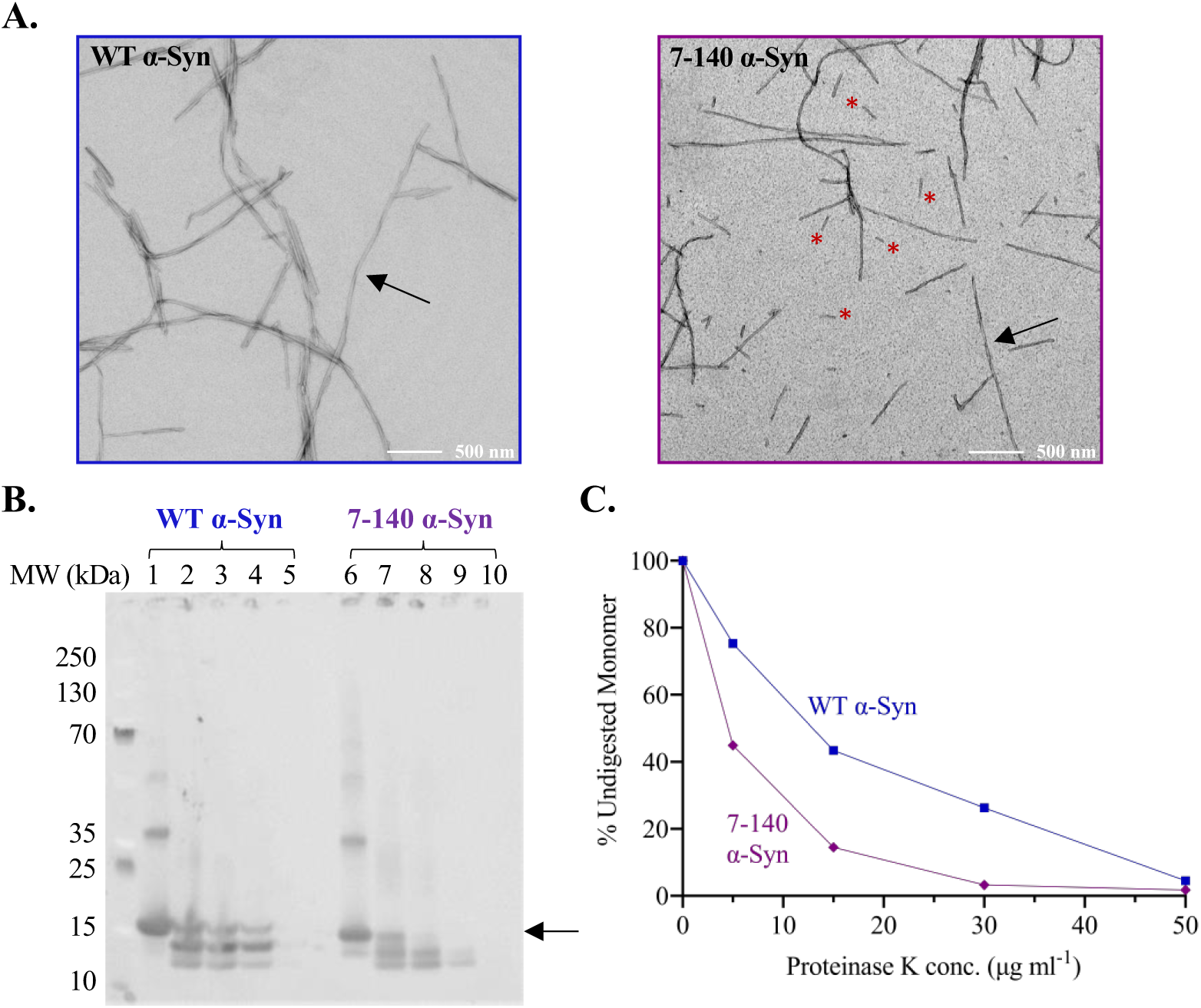
**A**. TEM images of WT and 7-140 α-syn fibrils after ∼ 140 h of aggregation. Arrows indicate fibrils with clear twisting, red asterisks highlight short fibrils. **B**. Proteinase K digestion of WT (lanes 1 – 5) and 7-140 (lanes 6 – 10) α-syn fibrils incubated with 0, 5, 15, 30 and 50 μg/ml proteinase K respectively. The arrow indicates the band that has been quantified and plotted **C**. The percentage of undigested monomeric WT (blue) and 7-140 (purple) α-syn at various protease concentrations.

### Concluding remarks

The aggregation of α-syn into amyloids is the hallmark of several neurodegenerative diseases including PD [6]. This process is highly complex because of the numerous α-syn PTMs, including N-terminal truncation [23, 26-28]. So far, the recombinant production of N-terminally truncated α-syn has been limited to protein variants starting with a Gly, retained after TEV cleavage, or a non-native Met for initiating translation. In this paper, we have described a facile method for the expression and purification of potentially any relevant N-terminally truncated fragment of α-syn without the need for the introduction of non-native residues or the use of a protease.

Our method allowed us to characterize the role of the first six N-terminal residues of α-syn in the amyloid aggregation of the protein. In our experimental conditions, we found that truncation of this region affects the aggregation of the protein and the stability of the amyloids (**Figure 3** and **4**). Our findings are in agreement with previous reports which found that the deletion of residues 2-11 of α-syn significantly delays aggregation [12], impairs membrane binding, and decreases toxicity in yeast [30]. We expect our method to be applied to generate additional N-terminally truncated α-syn variants to shed light into the mechanisms of the N-terminal region of α-syn.

## Conflict of Interest

*The authors declare that the research was conducted in the absence of any commercial or financial relationships that could be construed as a potential conflict of interest*.

## Author Contributions

RJT and DMV performed the experiments. All authors analyzed the data and wrote the paper. RJT and FAA conceptualized the work.

## Funding Sources

We thank UK Research and Innovation (Future Leaders Fellowship MR/S033947/1), the Alzheimer’s Society, UK (Grant 511), and Alzheimer’s Research UK (ARUK-PG2019B-020) for support. R.J.T. was supported by a scholarship by the Department of Chemistry (Imperial College London).

## Acknowledgements

The authors acknowledge Lisa Haigh and the Chemistry Mass Spectrometry facilities for ESI-MS experiments, and the Electron Microscopy Centre facilities at The Center of Structural Biology for TEM experiments.

## Abbreviations

α-Syn: α-Synuclein
PD: Parkinson’s disease
LB: Lewy Bodies
NAC: non-amyloid-β component
LLPS: liquid-liquid phase separation
PTM: post-translational modification
WT: wild-type
*E. coli*: *Escherichia coli*
TEV: tobacco etch virus
HEPES: 4-(2-hydroxyethyl)-1-piperazineethanesulfonic acid
EDTA: Ethylenediaminetetraacetic acid
CV: column volume
BME: β-mercaptoethanol
MBP: maltose binding protein
SEC: size-exclusion chromatography
SDS-PAGE: sodium dodecyl sulphate–polyacrylamide gel electrophoresis
ESI-MS: electrospray ionization mass spectrometry
ThT: Thioflavin T
CD: circular dichroism
MRE: mean residue ellipticity
*Sce*: *Saccharomyces cerevisiae*
VMA: vacuolar ATPase
CBD: chitin binding domain
TEM: transmission election microscopy

## Notes

### Competing Interest Statement

The authors have declared no competing interest.

