## Supplementary Material for "A Facile Method to Produce N-Terminally Truncated α-Synuclein"

### Supplementary Figures

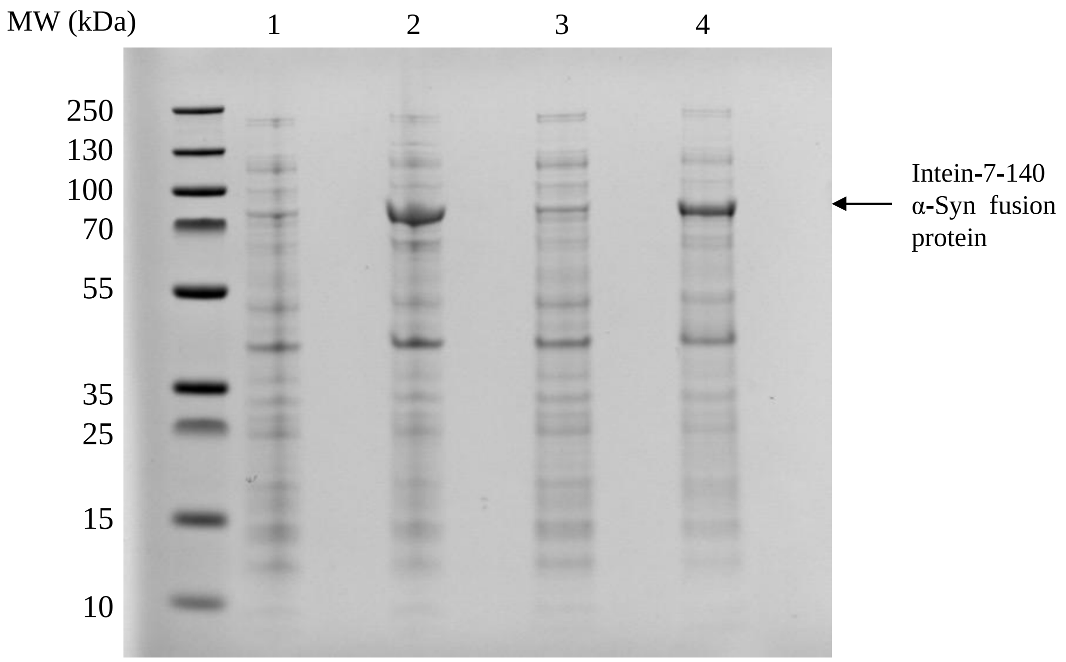


**Figure S1.** SDS-PAGE of cell lysate before and after induction of protein expression with 1 mM IPTG. We induced expression at 28 °C overnight (lane 2) and at 37 °C for 4 hours (lane 4). Lanes 1 and 3 contain the lysate of cell aliquots taken before induction at 28 °C and 37 °C respectively. The indicated band in lanes 2 and 4 corresponds to the 72.1 kDa intein-7-140 α-syn fusion protein.

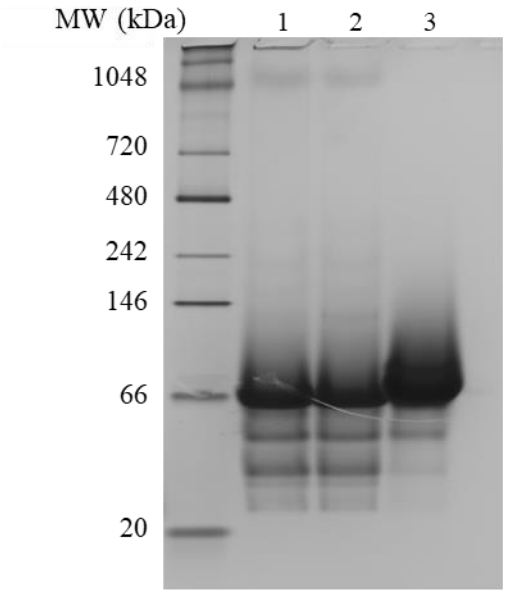


**Figure S2.** Native PAGE of 7-140 α-syn before (lane 2) and after (lane 1) boiling at 80 °C for 20 mins with conventionally purified WT α-syn (lane 3).

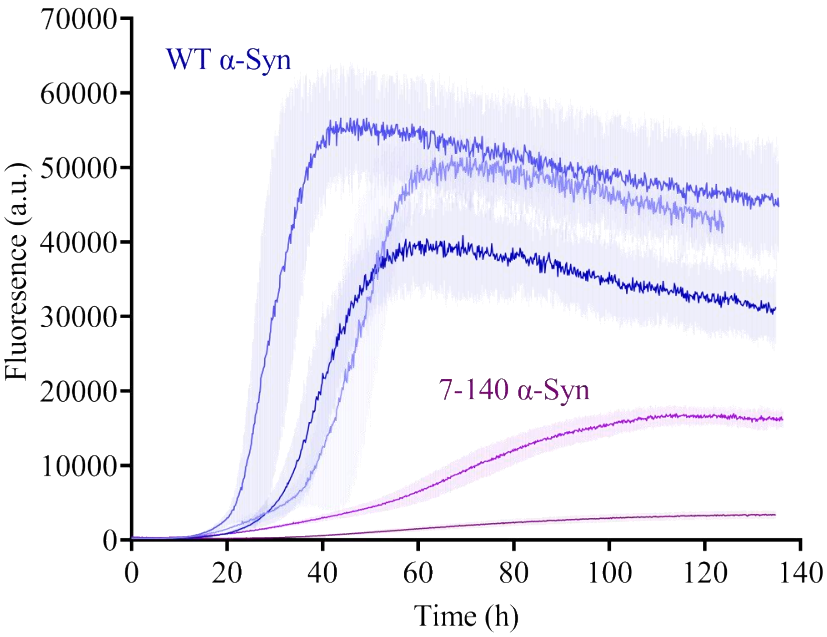


**Figure S3.** Raw data for the beaded aggregation of WT α-syn (blue) and 7-140 α-syn (purple). Error bars representing the standard deviation are shown as transparent bars. Each data set is the average of five repeats.

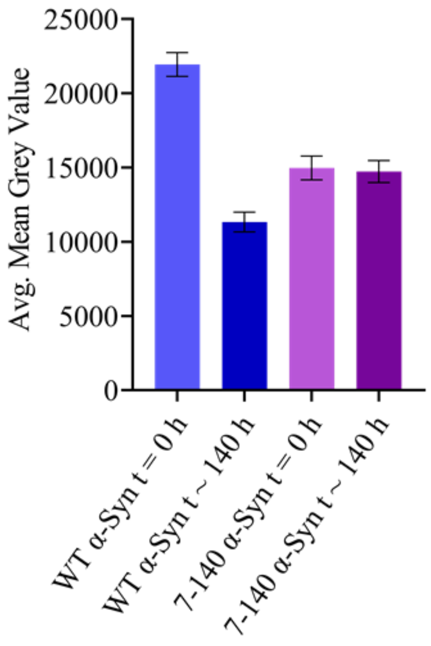


**Figure S4.** Mean grey values calculated using dot blot analysis of the soluble fraction before and after the aggregation of WT and 7-140 α-syn for ~ 140 h (n = 3, values represent means and error bars represent standard deviation).

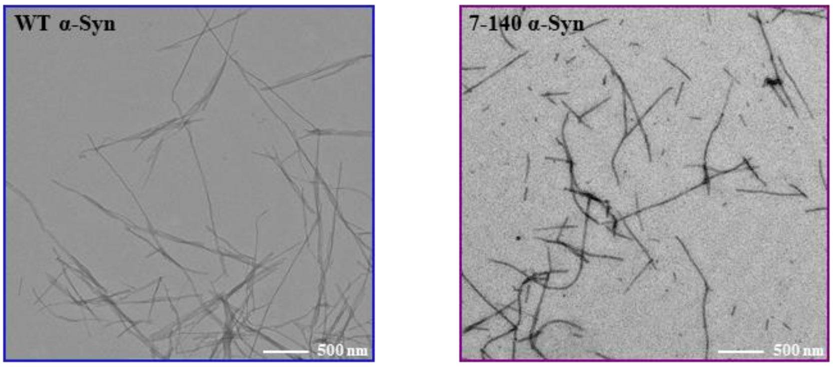


**Figure S5.** TEM images of WT and 7-140 α-syn fibrils after ~ 140 h of aggregation.
